# Learning Image Memorability with Feedback-Based Training

**DOI:** 10.1101/2023.12.18.572187

**Authors:** Cambria Revsine, Wilma A. Bainbridge

## Abstract

Memorability, or the likelihood that an image is later remembered, is an intrinsic stimulus property that is remarkably consistent across viewers. Despite this consistency in what people remember and forget, previous findings suggest a lack of consistency in what individuals subjectively believe to be memorable and forgettable. We aimed to improve the ability of participants to judge memorability using a feedback-based training paradigm containing face images (Experiment 1) or scene images (Experiment 2 and its replication and control experiments). Overall, participants were fairly accurate at categorizing the memorability of images. In response to the training, participants were able to improve their memorability judgments of scenes, but not faces. Those who used certain strategies to perform the task, namely relying on characteristic features of the scenes, showed greater learning. Although participants improved slightly over time, they never reached the level of ResMem, the leading DNN for estimating image memorability. These results suggest that with training, human participants can better their understanding of image memorability, but may be unable to access its full variance.

## Introduction

When we view a beautiful sunset, unique café, or striking face for the first time, we may think that we will remember it long after. But how accurate are people at predicting what they will remember versus what they will forget? And can individuals improve this skill in order to better understand their own memory capabilities?

Metacognition research has long attempted to answer these questions. Paradigms testing the metamemory phenomenon “judgments of learning” (JOLs) record participants’ predictions of their memory for certain items, before they perform a memory test on the same items (Rhodes, 2016). When comparing predictions to true memory performance, studies generally find moderate, above-chance accuracies of JOLs taken immediately after study (Arbuckle & Cuddy, 1969; Koriat, 1997; Mazzoni & Nelson, 1995; Nelson & Dunlosky, 1991). Metamemory theories suggest that individuals rely on various cues, including feelings of fluency and/or familiarity, to predict what they will remember (Koriat & Ma’ayan, 2005; Metcalf et al., 1993). However, these judgments are not always accurate, and salient perceptual features of an item, such as font size, have been shown to cause erroneous JOLs (Rhodes & Castel, 2008). Evidence on the effect of feedback on metamnemonic accuracy is mixed, with some studies finding that feedback does increase accuracy (Geurten & Meulemans, 2017; Koriat, 1997; Logan et al., 2012; McGillivray & Castel, 2011) but others finding a negative or null effect (Kornell & Rhodes, 2013; Sitzman et al., 2016).

However, the vast majority of JOL research has used verbal materials, with only a handful of studies testing metamemory for naturalistic images (Kao et al., 2005; Undorf & Bröder, 2021; Saito et al., 2022). Although findings suggest that there may be similarities in the basis and accuracy of JOLs for both types of stimuli (Undorf & Bröder, 2021), work on the “picture superiority effect” implies an inherent difference in that pictures are better remembered than words (Hockley, 2008; Paivio & Csapo, 1973). Furthermore, to our knowledge, no study has investigated the effect of feedback on metamemory for images. This lack of research is surprising, given the recent surge in interest for image memory due to the emergence of an “objective memory measure” for images, dubbed memorability. Memorability, or the likelihood of a stimulus to be remembered, is an intrinsic image property that is highly consistent across viewers—i.e., people tend to remember and forget the same images as one another (Bainbridge et al., 2013; Isola et al., 2011). The consistency of this property has been shown across a range of stimulus types (Bainbridge, 2017; Isola et al., 2014; Ongchoco et al., 2022; Xie et al., 2020) and task contexts (Broers et al., 2018; Bylinskii et al., 2015; Goetschalckx et al., 2018). The determinants of memorability are still largely unknown; however, studies have demonstrated a greater influence of semantic over low-level visual features (Bainbridge et al., 2013; Isola et al., 2014; Kramer et al., 2023).

Critically, a memorability angle can offer new insights into metacognitive questions of how, and how well, individuals can predict the images they will remember and forget. With this approach, instead of judging what they themselves will remember, participants judge what others would tend to remember, i.e., *subjective memorability*. Prior work using such an approach has suggested that, contrary to the consistency in objective memorability across people, there is no consistency in subjective memorability ratings (Bainbridge, 2017). Furthermore, when participants rated scene photographs on whether they thought the images were memorable, researchers found no correlation with true memorability, implying that people have poor ability to predict the image property (Isola et al., 2014). In contrast to these findings, a recent study testing both JOLs and subjective memorability judgments of face and object images found that participants’ judgments were predictive of both measures, though with some remaining unexplained variance (Saito et al., 2022). The current study aims to arbitrate between these two accounts utilizing stimulus categories used by both sides of the debate. Importantly, however, although evidence is mixed as to the initial accuracy of these judgments, there is room to improve this accuracy.

### Although it is unclear whether humans have insight into memorability, surprisingly, deep neural network (DNN) models have impressive performance at making those same judgments

ResMem is currently the top-performing DNN for estimating image memorability, achieving a 0.67 correlation with true hit rates (Needell & Bainbridge, 2022). In addition, a recent study found that ResMem captures aspects of memorability that are seemingly inaccessible to humans (Zhao et al., 2023). Given that a DNN can perform this task, we wondered if human participants receiving feedback could likewise improve their imperfect understanding of memorability, perhaps by learning the additional features ResMem uses. It is possible that this image property is simply too opaque a concept for humans to estimate. This idea is supported by the fact that regression models predicting memorability from human-labeled image features have only been able to explain 50% of its variance (Bainbridge et al., 2013); therefore, the determinants of memorability are still largely a mystery. Alternatively, with effective training, participants may learn to distinguish memorable and forgettable images. Recent findings showing that famous artworks are inherently more memorable support this possibility, as this suggests that successful artists can discern memorability (Davis & Bainbridge, 2023). This outcome would not only shed light on how humans generate and improve their metacognitive judgments for pictures, but could provide further insight into what makes an image memorable.

To test these questions, we ran multiple experiments in which participants viewed a sequence of highly memorable and forgettable face images (Experiment 1) or scene images (Experiment 2 and its replication and control experiments), categorized the memorability of each image, and received the correct answer following each trial. We aimed to see whether this training paradigm could improve participants’ memorability judgments. In sum, we found that subjective memorability ratings generally aligned with true ratings, but failed to reach DNN-level performance. Participants exhibited a slight but significant improvement in classifying the memorability of scene images over time, but showed no improvement for face images. Certain participant-reported strategies that focused on the characteristic features of scenes proved most effective for learning memorability. These results provide support to the idea that individuals can improve upon their initial sense of what is memorable, yet emphasize the complexities of memorability that limit this effect.

## Methods

### Experiment 1: Face Memorability Training Experiment

#### Participants

Participants were recruited through Prolific, an online platform for researchers to conduct behavioral experiments. All participants were prescreened for U.S. nationality and fluency in English. An approval rating of ≥ 90% and completion of ≥ 50 previous experiments on the site were required to participate in the study. 100 participants successfully completed the task (18 – 72 years old, mean age = 34.60, SD = 13.62). Compensation was provided at a rate of $6.50 per hour. Participants provided electronic informed consent prior to the experiment, following the University of Chicago Institutional Review Board guidelines (IRB19-1395).

We determined our sample size from previous work finding that a stable measure of memorability is reached at 80 participant ratings per image (Isola et al., 2014), and recruited an additional 20 participants to account for the noisier data found in online studies. As in prior online memorability studies (Bainbridge et al., 2013; Davis & Bainbridge, 2023; Isola et al 2014; Kramer et al., 2023) we did not implement an age limit in any of the present experiments. However, we ensured that results replicated after removing participants ≥ 65 years old from the analyses (3 participants in Exp. 1 and 5 participants in Exp. 2; **Supplementary Text 1**).

#### Stimuli

Face images were used as the stimuli for Experiment 1. Images came from the 10k US Adult Faces Database, a large-scale photograph database of more than 10,000 adult individuals closely matching the demographics of the U.S. population (Bainbridge et al., 2013). These images are cropped with an oval to exclude background area, and measure 256 pixels in height. A subset of 2,222 images in the database have corresponding memorability ratings, including hit rate (HR) and false alarm rate (FAR), both of which were calculated by averaging memory performance from ∼80 participants per image from a previous behavioral experiment (Bainbridge et al., 2013). In the current study, we define objective memorability as image HR, however we also use FAR in analysis. Face images were chosen for this experiment as they have been used extensively to characterize objective memorability (Bainbridge et al., 2013; Bainbridge, 2017), and have been shown to elicit accurate subjective memorability judgments (Saito et al., 2022). Faces also serve as an interesting stimulus set because their HRs show high variance with a relatively low mean (mean HR = 0.52, SD = 0.13; Bainbridge et al., 2013), so that there is a large memory performance difference between memorable and forgettable images that could make it easier to learn the difference between memorable and forgettable sets. Further, as faces are from the same base-level image category with similar semantic and visual features, participants are less able to rely on a singular type of feature to perform the task.

Stimuli chosen for this experiment consisted of 90 highly memorable images and 90 highly forgettable images, sampled from the upper and lower ends of the HR distribution of the face database, respectively, for a total of 180 images. We chose to use highly memorable and forgettable stimuli in both experiments, in order to test whether participants would show a learning effect under these relatively easy task conditions. Memorable and forgettable image sets were matched for age, race, and gender, to ensure that there were no overall differences in demographics between the two stimulus conditions. This resulted in a memorable image set with HRs ranging from 0.6 to 0.86 (HR ≥ 0.75*SD above the database mean), and a forgettable image set with HRs 0.15 to 0.42 (HR ≤ 0.75*SD below the mean). In addition to memorability ratings, images in the 10k US Adult Faces Database have crowdsourced ratings of 20 psychology and memory-related attributes and their antonyms (e.g., sociable, aggressive, familiar) (Bainbridge et al., 2013). These attribute ratings were used in analysis.

#### Procedure

Prior to beginning the task, participants were informed that they would see many different images, each of which was either a highly memorable or highly forgettable image. Memorable images were defined in the instructions as “consistently remembered much more frequently by participants in previous studies” than forgettable images. During the experiment, each trial began with a central fixation cross presented for 500 ms, followed by a single face image at the center of the screen (**Figure 1**). Participants freely viewed the face, and then indicated with a key press whether they believed it to be memorable or forgettable, by pressing the *J* or *F* key, respectively. There was no time limit for this stage; the image remained on the screen for as long as participants took to answer. Immediately following their response, feedback appeared below the image indicating whether the trial was answered correctly or incorrectly, along with the true memorability category of the stimulus. This screen was displayed for three seconds, to allow participants sufficient time to study the image along with its memorability category. Following the reappearance of the fixation cross, a new image was presented at the center of the screen. The task continued in this manner with a new face image each trial, for a total of 180 trials. All participants saw a randomized order of the same 180 images. The task also included 10 evenly spaced attention check trials, in which participants were shown a face that did not appear elsewhere in the task and had to indicate the face’s gender. Participants who did not correctly answer at least 60% of these trials (6% of total participants in Exp. 1) were excluded from analysis and replaced with new participants. In addition, four break points were interspersed throughout the experiment. In total, participants took about 20 minutes to complete the task.

**Figure 1.**
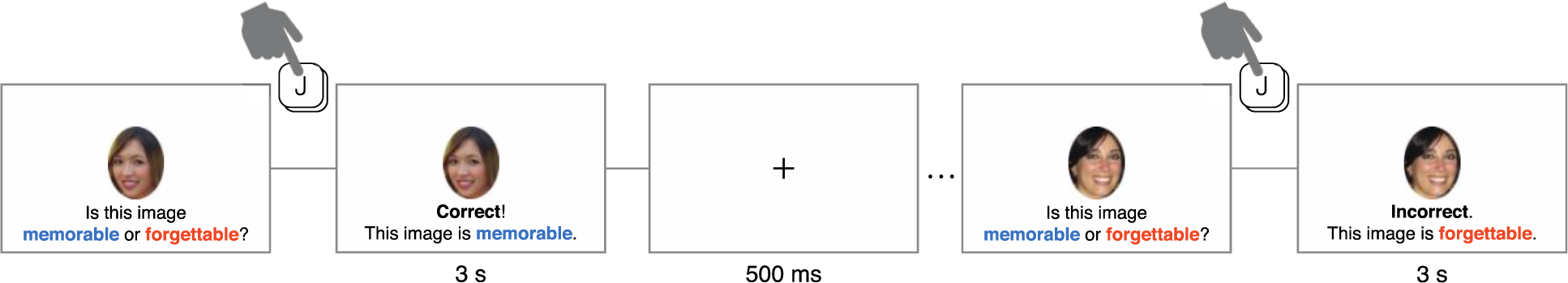
The Memorability Judgment Task. *Note.* In Experiment 1, participants viewed a sequence of face images, and for each image indicated with a button press whether they thought it was a memorable face (by pressing the *J* key) or a forgettable face (by pressing the *F* key). Immediately following their response, feedback was displayed below the image informing the participant of the image’s correct memorability category. This was repeated for 180 unique images. The design of Experiment 2 was identical to that of Experiment 1, except that stimuli consisted of scene images.

Following their completion of the task, participants were asked to provide written responses to three questions. These included: “What strategies did you use to perform this task?”, “What do you think was the difference between memorable and forgettable images?”, and “How, if at all, did your ideas about this difference change over the course of the experiment?”

### Experiment 2: Scene Memorability Training Experiment

#### Participants

A separate group of 100 participants on Prolific were recruited to participate in a second experiment (18 – 82 years old, M = 34.73, SD = 14.99). 47% of participants were female, 49% were male, and 4% marked “other” or preferred not to answer. All recruitment and screening procedures were identical to those used in Experiment 1.

#### Stimuli

Scene images were used as the stimuli for Experiment 2 as, like faces, they have often been used to characterize memorability (Isola et al., 2011, 2014), but differ from faces in their semantic and perceptual variability. Further, prior work has shown negative, non-significant correlations between objective and subjective memorability ratings for scenes (Isola et al., 2014), suggesting that intuitive judgments begin as very inaccurate. Images used here came from the Scene Understanding (SUN) database (Xiao et al., 2010). This database contains more than 100,000 natural images of places and scenes across hundreds of scene categories (e.g., kitchen, river). A subset of 2,222 SUN images have corresponding HR and FAR scores, as computed by a previous behavioral experiment (Isola et al., 2011). Compared with the face database in Exp. 1, these scene images have a relatively high mean HR, with comparable variance (Mean HR = 0.68, SD = 0.14). As in Exp. 1, 90 highly memorable and 90 highly forgettable images were pseudorandomly selected from the extremes of the HR distribution of this database, for a total of 180 stimuli. These stimuli excluded any scenes containing people, as they have been shown to be more memorable than images without people (Isola et al., 2011). Memorable and forgettable image sets were matched for scene category whenever possible, and were counterbalanced for number of indoor/outdoor scenes and manmade/natural scenes. This resulted in a memorable image set with HRs ranging from 0.81 to 0.95 (HR ≥ 1*SD above the database mean), and a forgettable set with HRs 0.19 to 0.54 (HR ≤ 1*SD below the mean). All stimuli were cropped to 256 x 256 pixels for use in this experiment.

#### Procedure

The design of Experiment 2 was identical to that of Experiment 1, with participants instead judging the memorability of scene stimuli. During the attention check trials, participants were shown a scene image that did not appear elsewhere in the task, and indicated whether the image was of an indoor or outdoor scene. 5% of total participants in Exp. 2 failed this attention check, and were replaced with new participants.

### Replication and Control Experiments

A pair of follow-up experiments were run, to compare with the results of Experiment 2. We ran a replication experiment with a separate group of 100 participants from Prolific (19 – 74 years old, M = 39.16, SD = 14.48; 50% male). The procedure of this experiment was identical to that of Exp. 2, except with a stricter attention check criterion (80%). Along with this experiment, we ran a no-feedback control experiment with another group of 100 Prolific participants (19 – 77 years old, M = 41.28, SD = 13.80; 51% male). The procedure of this experiment was also similar to that of Exp. 2, except that following each trial, participants were simply reminded of their own response instead of receiving feedback on the correct image category. Here, the attention check criterion was also set at 80%.

### Analyses

#### Subjective memorability ratings

To obtain a measure of subjective memorability analogous to hit rate (objective memorability), we calculated the proportion of responses to each image where participants guessed the image was memorable (versus forgettable), which we call the “guessed memorability rating”. We also calculated the average accuracy for each image. For memorable images, this was equal to the guessed memorability rating, while for forgettable images, this was equal to 1 – the guessed memorability rating.

#### Split-half consistency analysis

To measure the agreement between participants in terms of their memorability judgments, we ran a split-half consistency analysis on guessed ratings, as described in Isola et al. (2011) and Bainbridge et al. (2013). In short, the participant pool was randomly split into two equal groups, and average guessed memorability ratings were calculated for all images in each group. A Spearman rank correlation was then measured between groups. Chance level consistency was also calculated by correlating group 1’s ranking with a shuffled group 2’s ranking. This was repeated 1000 times, with the resulting split-half correlation values averaged to produce the final consistency value. The significance of this value was determined using a nonparametric permutation test, comparing the average correlation value with the 1000 shuffled correlations. A high average correlation value indicates that participants generally agreed on the images they judged as memorable and forgettable.

#### Within-participant linear regression

To model learning over time for each participant, we split trials into 10 time bins of 18 trials each, and calculated each participant’s average accuracy in all 10 bins. We then logit-transformed the accuracy percentages, to account for their bounded nature, before fitting a simple linear regression predicting each participant’s accuracy over time. To test for overall significance, a one-sample t-test was conducted to compare the mean of all 100 regression coefficients to zero. This provided a general measure of how well participants were improving overall.

#### Signal detection measures of bias and discriminability

To investigate classification performance beyond just changes in average accuracy, we calculated signal detection measures of bias and discriminability. As a measure of discriminability between stimulus conditions, *d’* was computed as *norminv(Hit Rate) - norminv(False Alarm Rate).* We calculated this measure for each participant, separately for the first and second half of the task. A paired t-test was then run to test whether *d’* values changed significantly from half 1 to half 2. As a measure of bias, criterion values were computed as - *0.5*(norminv(Hit Rate) + norminv(False Alarm Rate))*. As with *d’*, criterion values were calculated for half 1 and 2 for each participant, and a paired t-test was used to measure a change in criterion values.

#### Mixed-effects logistic regression model

We investigated the presence of more fine-grained learning using a mixed-effects logistic regression model to predict the binary outcome of each trial—either correct or incorrect—across participants. The formula used to specify the model was:

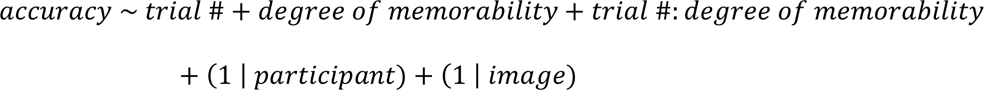

Thus, the model included two predictor variables as fixed effects: 1) *trial number* and 2) *degree of memorability*, or how memorable or forgettable an image was, calculated as abs(Hit Rate – 0.5). This predictor, which results in higher values the closer stimulus hit rate is to 0 or 1, was included in the model as we hypothesized that more extremely memorable and forgettable images might be more easily identified as such by participants. We also included an interaction term between trial number and degree of memorability. Intercepts for participant and image were included as random effects. Both predictors were z-scored prior to fitting the model. We specified a binomial distribution, logit link function, and maximum likelihood as the method for estimating model parameters.

#### Individual differences analysis

To examine individual differences in learning, we categorized participants by their responses to the questions asked following the task regarding their strategies for judging memorability. These categories were determined according to common themes identified across responses that applied to at least 10 participants each. Categories in Experiment 1 (faces) included 1) distinguishing facial feature(s) (e.g., eyes, smile), 2) general face traits (e.g., symmetry, attractiveness), 3) expression/emotion, 4) intuition/gut feeling, 5) image quality/lighting, 6) whether the participant thought they would remember the face, 7) whether the image caught their attention, and 8) consciously incorporating the task feedback. Categories in Experiment 2 (scenes) included 1) specific objects or places (e.g., landmarks, signs), 2) general scene traits (e.g., aesthetics, simplicity), 3) color, 4) intuition/gut feeling, 5) image quality/lighting, 6) whether the participant thought they would remember the scene, and 7) consciously incorporating the task feedback. Participants could be assigned to more than one response category; thus, we also analyzed combinations of two categories with at least 10 participants each. In each individual and joint category group, we tested for a significant change in performance using the within-participant linear regression analysis described above, with a false discovery rate (FDR) procedure to correct for multiple comparisons (*q* < 0.05).

#### Nonparametric permutation tests

We used a nonparametric permutation test anytime we directly compared two correlation coefficients in the results. Pairs of x- and y-values were randomly shuffled across both data sets before computing the correlations on the shuffled data, and taking the difference between the correlation coefficients. We did this over 10,000 iterations, resulting in a null distribution of correlation differences. The proportion of this null distribution more extreme than the true correlation difference determined the significance of the difference between the two correlations.

### Transparency and Openness

We report how we determined our sample size, all data exclusion criteria, all manipulations, and all measures in the study. All data are available at https://osf.io/34v26/ (data will be made available upon publication of the paper). Data were analyzed using MATLAB, version 9.12.0.1956245 (R2022a). This study’s design and its analysis were not pre-registered.

## Results

### Accuracy and consistency of guessed memorability ratings

In this study, participants categorized the memorability of highly memorable and forgettable face or scene images during a feedback-based training paradigm. In Experiment 1 (faces), average across-subject accuracy was 63.7% (SD=7.4%). In Experiment 2 (scenes), average accuracy was 58.3% (SD=7.2%). There was no difference in average response time between Experiment 1 and 2 (*p*>0.05), nor between memorable and forgettable stimuli in either experiment (both *p*>0.05).

To obtain a measure of average performance by image, we next compared participants’ memorability judgments with true stimulus HRs. There was a high correlation between guessed and true memorability ratings for faces (ρ=0.72, *p*<0.001), and a moderate correlation for scenes (ρ=0.42, *p*<0.001) (**Figure 2a**). The correlation for faces was significantly greater than for scenes, confirmed by a permutation test over 10,000 iterations (*p*<0.001). Thus, participants were generally accurate at judging stimulus memorability, more so for faces than scenes. We tested the relationship between guessed memorability ratings and face attribute ratings of the Exp. 1 stimuli, and found that no correlation was greater than that of guessed and true memorability (all *p*>0.05, confirmed by 10,000-iteration permutation test), suggesting that participants did not rely completely on a specific attribute when judging memorability (**Supplementary Table 1**). Results from a split-half correlation analysis revealed high agreement across participants in memorability judgments for both faces (ρ=0.81, *p*<0.001) and scenes (ρ=0.87, *p*<0.001) (**Supplementary Figure 1**). **Figure 3** shows example scene images that were categorized correctly and incorrectly by participants.

**Figure 2.**
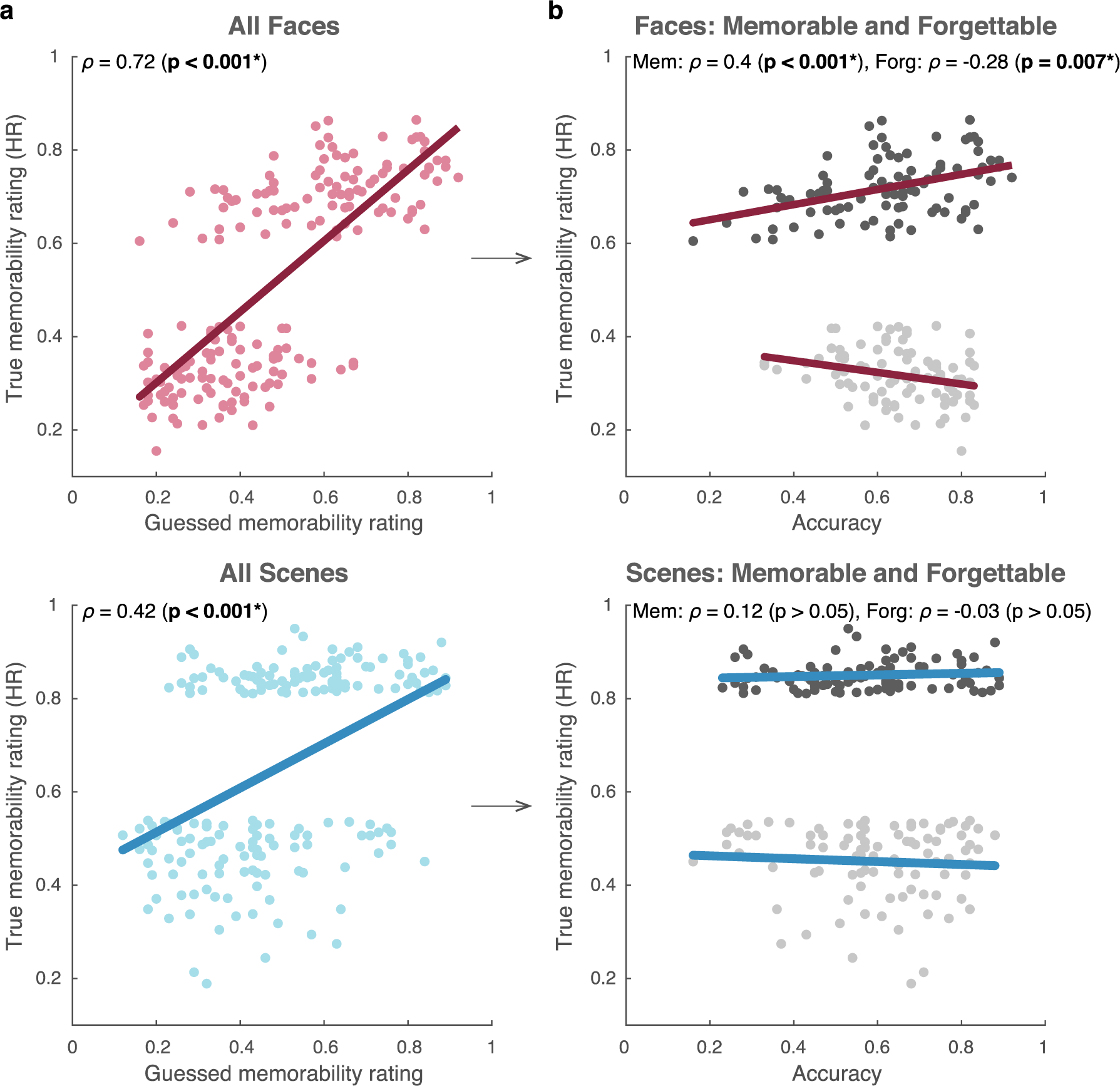
Relationship Between Participant Judgments and True Memorability Ratings. *Note.* a) Average guessed memorability rating is plotted along the x-axis and true HR is plotted along the y-axis for individual stimuli, represented by points in the scatterplots. The correlation between guessed and true HR was significant for both face images (*top*) and scene images (*bottom*). b) As in a), true HR is plotted along the y-axis. Average accuracy for each image is plotted along the x-axis. For memorable stimuli (dark gray), this measure equals the guessed memorability rating in a). For forgettable stimuli (light gray), this equals 1 – the guessed memorability rating. For faces (*top*), the correlation between accuracy and true HR was positive for memorable stimuli and negative for forgettable stimuli. For scenes (*bottom*), this correlation was non-significant for both memorable and forgettable stimuli. All lines correspond to linear least squares fit.

**Figure 3.**
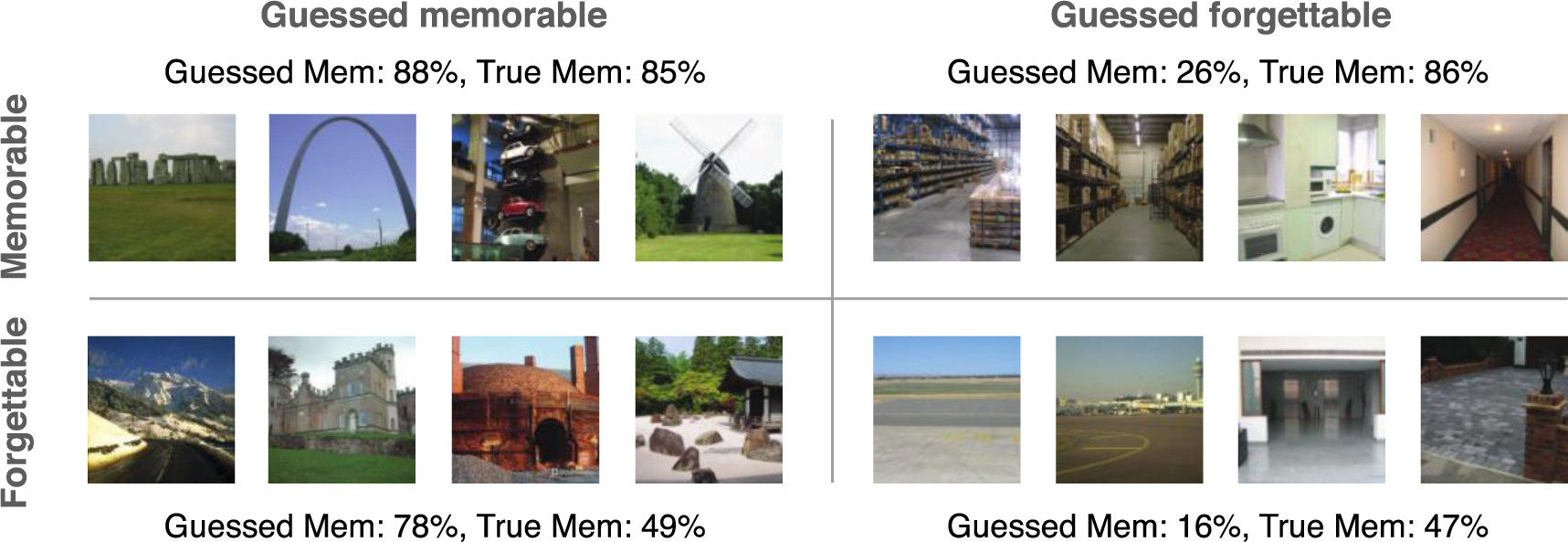
Examples of Correctly and Incorrectly Categorized Scene Stimuli. *Note.* The four quadrants include correctly identified memorable (*top left*) and forgettable (*bottom right*) stimuli, as well as forgettable stimuli that were incorrectly judged as memorable (*bottom left*) and memorable stimuli that were incorrectly judged as forgettable (*top right*). The first value by each quadrant corresponds to the mean guessed memorability rating of the four images, followed by their mean true memorability rating.

We next examined this relationship between HR and average accuracy, separately for memorable and forgettable stimuli. For face stimuli, memorable faces exhibited a positive correlation between accuracy and HR (ρ=0.40, *p*<0.001) and forgettable faces exhibited a negative correlation (ρ=-0.28, *p*=0.007) (**Figure 2b**). Thus, the more a face was memorable or forgettable, the more accurate participants were at determining its memorability. For scene stimuli, however, we found no such significant correlations for either memorable (ρ=0.12, *p*>0.05) or forgettable scenes (ρ=-0.03, *p*>0.05).

To test whether stimulus False Alarm Rates (FAR) also influenced subjective memorability judgments, we compared these two measures as well. We observed a negative correlation for faces (ρ=-0.25, *p*<0.001), but no significant correlation for scenes (*p*>0.05) (**Supplementary Figure 2**). Thus, participants did not fall for the false familiarity of high FAR images when making memorability judgments.

### Learning memorability over time

We addressed our central question of whether participants learned memorability by analyzing their performance over 10 equal time bins. For faces, average across-subject accuracy was 61.2% (SD=13.6%) in the first time bin and 64.3% (SD=12.3%) in the last time bin. For scenes, average accuracy was 56.7% (SD=12.4%) in the first bin and 59.4% (SD=15.9%) in the last bin. Performance was significantly higher for faces than scenes in all but one time bin (two-sample *t*-tests: *p*<0.05, FDR-corrected *q*<0.05; bin 7: *p*>0.05; **Supplementary Table 2**).

As an initial look into the effect of training over time, we then fit a linear regression for each participant, predicting logit-transformed accuracy across the 10 time bins. In comparing the resulting regression coefficients to zero, we observed no significant difference for faces (mean beta=0.008, SD=0.062); *t*(99)=1.23, *p*>0.05 (**Figure 4a**). For scenes, betas were significantly greater than zero (M=0.016, SD=0.065); *t*(99)=2.53, *p*=0.013 (**Figure 4b**). We also compared learning in the first and second half of trials directly, and found significant improvement in the first half of Exp. 1 (M=0.037, SD=0.17); *t*(99)=2.14, *p*=0.034, but no difference between halves (*p*>0.05; **Supplementary Figure 3**). Thus, participants showed slight improvement in their ability to determine scene memorability, but not face memorability, over the full task.

**Figure 4.**
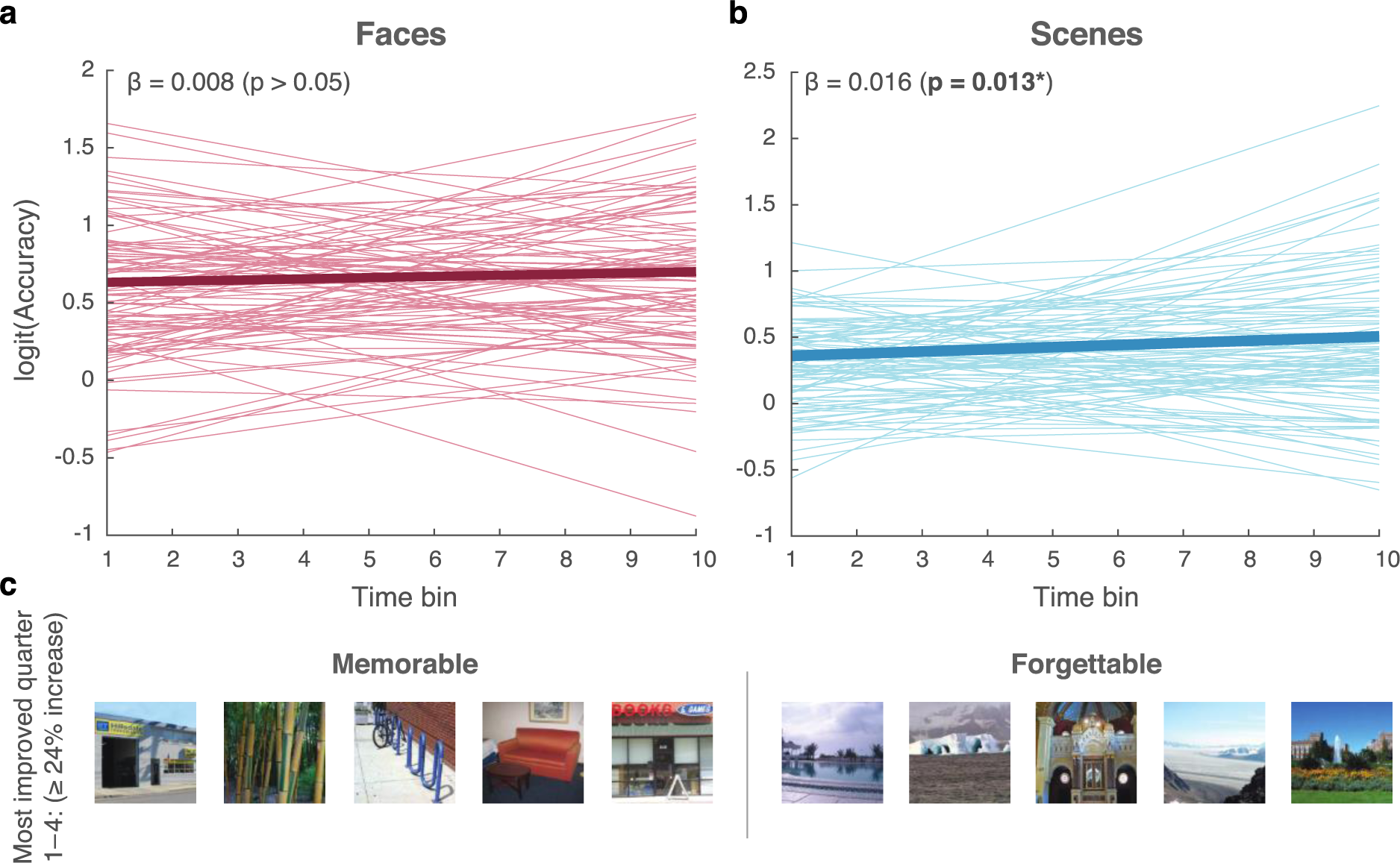
Task Performance Over Time, and Most Improved Stimuli. *Note.* a-b) Each line corresponds to a simple linear regression predicting logit-transformed average accuracy (y-axis) from time (x-axis) for a single participant. Bolded lines correspond to the average regression lines. a) For faces, regression slopes were not significantly different from zero. b) For scenes, slopes were significantly greater than zero. c) Memorable and forgettable scene stimuli with the greatest average improvement in accuracy when viewed in the first quarter compared with the last quarter of trials. All stimuli shown here increased in accuracy by at least 24%.

We then examined whether the observed learning effect in Experiment 2 replicated in an additional experiment, using the same analysis approach. We chose just to replicate Exp. 2, as participants in Exp. 1 showed little evidence of learning. As in Exp. 2, participants exhibited a significant increase in accuracy over time (M=0.026, SD=0.067); *t*(99)=3.93, *p*<0.001 (**Supplementary Figure 4**). Next, we analyzed results from a no-feedback control experiment, to test whether improved performance over time was indeed due to learning from feedback. Here, participants showed no change in accuracy (M=-0.004, SD=0.066); *t*(99)=-0.615, *p*>0.05. There was also a significant difference between regression coefficients in the replication and control experiments (*t*(99)=3.02, *p*=0.003). These results suggest that the feedback-based training did improve participants’ subjective memorability judgments for scenes, whereas they did not improve in the absence of feedback.

To ensure that this improvement was due to an increase in discriminability between stimulus conditions and not a decrease in response bias, we measured *d’* and criterion values of all participants, and compared these values in the first and second half of trials. For faces, we did not observe a significant change from half 1 to half 2 for either *d’* (half 1: M=0.73, SD=0.44, half 2: M=0.78, SD=0.49); *t*(99)=1.16, *p*>0.05, or criterion values (half 1: M=0.07, SD=0.26, half 2: M=0.03, SD=0.37); *t*(99)=-1.11, *p*>0.05. For scenes, *d’* significantly increased from half 1 (M=0.39, SD=0.38) to half 2 (M=0.50, SD=0.49); *t*(99)=2.73, *p*=0.008, but criterion values did not change (half 1: M=0.05, SD=0.29, half 2: M=0.01, SD=0.33); *t*(99)=-1.34, *p*>0.05. Importantly, this suggests that the improvement observed in Exp. 2 was not due to a decrease in the bias of participants’ judgments, but rather an increase in participants’ discriminability of memorable and forgettable stimuli.

Next, we fit a mixed-effects logistic regression model to investigate the trial-by-trial effects of time and memorability on learning, across participants. Here, we predicted the outcome of each trial (correct or incorrect) from trial number and degree of memorability—how memorable or forgettable an image was. In predicting accuracy in Experiment 1, our model had an adjusted *R^2^*of 0.11. Degree of memorability was predictive of accuracy for faces (*β*=0.23, *p*<0.001), mirroring our earlier finding, but trial number was non-significant (*β*=0.02, *p*>0.05) (**Supplementary Table 3**). The interaction term between predictors was also non-significant (*β*=-0.01, *p*>0.05). In predicting accuracy for scenes, our model had an adjusted *R^2^* of 0.15. In contrast to Exp. 1, however, trial number was significantly predictive (*β*=0.04, *p*=0.018), but degree of memorability was not (*β*=-0.02, *p*>0.05). The interaction term was marginally significant (*β*=0.03, *p*=0.054). This interaction appears to reflect trial number positively affecting accuracy more for stimuli with high degrees of memorability, i.e., the most memorable and forgettable images (**Supplementary Table 4**). These results echo our previous results suggesting differences in participants’ learning dependent on stimulus category. While degree of memorability largely drove accuracy for faces, trial number significantly affected accuracy for scenes, supporting the idea that participants learned their memorability over time.

### Comparison with DNN performance

We next wondered how the performance of the human participants in our study compared to that of ResMem, a DNN trained to estimate image memorability (Needell & Bainbridge, 2022). Testing ResMem on the same scene stimuli used in Experiment 2 resulted in a 0.69 correlation between estimated and true HRs (*p*<0.001), consistent with previously reported performance levels. This value was significantly greater than the 0.42 correlation reached by participants in our study (*p*=0.004; confirmed by 10,000-iteration permutation test). Because ResMem was trained on scene images, we did not use it to predict memorability of the face images in Experiment 1.

A more accurate comparison to ResMem’s performance is the final performance level reached by participants. To estimate this, we performed the same correlation analysis within the last quarter of trials (trial 135-180; **Figure 4c**). For scenes, this “final” correlation between guessed and true memorability was 0.46 (*p*<0.001) (**Supplementary Figure 5**), still significantly below that of ResMem (*p*=0.01; confirmed by 10,000-iteration permutation test).

### Individual differences in learning

Within our participant pool, did some individuals learn memorability better than others? To answer this, we grouped participants according to their responses to the post-task questions regarding strategies and ideas about memorability (**Supplementary Table 5)**. Grouping participants by strategy resulted in only one response category showing significant learning. Participants in Experiment 2 who used specific object(s) or place(s) to determine scene memorability exhibited an increase in performance (*t*(62)=3.34, *p*=0.001, FDR-corrected *q*<0.05). Participants who used this strategy in conjunction with the colors present in the scene also significantly improved (*t*(17)=2.94, *p*=0.009, FDR-corrected *q*<0.05). All other individual and joint strategies resulted in either marginal improvement (failing FDR correction) or no change in performance (**Supplementary Table 6**). These findings illustrate differences in how participants learned, or did not learn, the memorability of stimuli, as a result of their own approaches to the task. The most effective strategy, focusing on specific places or objects in the images, suggests that characteristic features of scenes may strongly influence their memorability, more than more general and/or low-level image features.

## Discussion

In this study, we aimed to determine whether people can improve their imperfect understanding of image memorability, by examining participants’ memorability judgments over a repetitive training paradigm. Participants exhibited slight but significant and replicable learning over time in response to scene images, but not in response to face images; in fact, memorability judgments of faces depended more on how memorable or forgettable the images were. Thus, we found limited evidence for the ability of individuals to learn memorability, dependent on stimulus category.

This work is the first to our knowledge to attempt to improve individuals’ image memorability judgments. However, it parallels previous findings testing both the accuracy and consistency of subjective memorability judgments in the absence of training. We found that judgments were consistent across participants and were moderately predictive of objective memorability, similar to the findings by Saito and colleagues (2022) for face and object images. In contrast, our results differ from those of Isola et al. (2014) who found no correlation between subjective and objective memorability scores for scenes, and Bainbridge (2017) who reported no consistency in the faces participants rated as memorable. One possible explanation for this difference could be the differences in stimulus selection across these studies. While the previous two studies used images spanning a wide range of memorability, we tested participants only on images with very high and low HRs, which are likely easier to categorize, as suggested by the results in Experiment 1. The fact that our study involved explicit training is another potential reason for these inconsistent findings, as training slightly improved the accuracy of memorability judgments in Exp. 2.

Though we tested participants’ subjective memorability judgments, our findings contribute to the related metamemory literature, providing additional evidence that judgments of learning are generally accurate (Arbuckle & Cuddy, 1969; Rhodes, 2016; Undorf & Bröder, 2021). This is especially informative as JOL research until recently has disproportionately focused on verbal materials; thus, our results suggest similar accuracy of JOLs for pictures. Our work is also the first to address the effect of feedback on JOLs for images. As with previous studies investigating its effect on JOLs for verbal materials (Koriat, 1997; Kornell & Rhodes, 2013), we found mixed evidence for the ability of feedback to improve subjective memorability judgments. We also failed to find strong evidence for classic theories of metamemory, suggesting potential differences in the basis of judgments for images and words. For example, the cue-familiarity hypothesis states that individuals use feelings of familiarity to predict what they will remember (Metcalf et al., 1993). However, we found no evidence for this idea in our data—in fact, participants in Exp. 1 predicted images with higher false alarm rates, representing familiarity, to be less memorable. This result aligns with the “mirror effect” of recognition memory, in which items with higher hit rates tend to have lower false alarm rates (Wixted, 1992). Another recent theory suggests that salient perceptual features of an item, such as font size, may falsely influence JOLs (Rhodes & Castel, 2008). Consistent with this theory, it is possible that the feedback participants received in the current study may have pointed them to parts of the images that contribute more to memorability, or conversely, misdirected them to less relevant features. Results of the individual differences analysis support this notion; participants who used features or objects denoting a scene’s conceptual information, which previous studies have found to be strongly related to memorability (Kramer et al., 2023; Needell & Bainbridge, 2022), significantly improved their judgments. In contrast, participants who relied on properties like the aesthetics or color of scenes, both of which are only weakly correlated with image memorability (Isola et al., 2014; Dubey et al., 2015), did not show significant learning. Although we identified some parallels with previous metamemory work, the remaining differences highlight the complex nature of image memorability, as well as the need for more research on the determinants of this property.

Adding to this complexity, we observed interesting differences within our results dependent on stimulus category, whereby participants were more accurate overall at judging face memorability, but only improved from feedback in response to scene images. A potential explanation for this discrepancy is the fact that faces comprise a single category and are thus relatively visually homogeneous, while scenes include many subcategories and can be very perceptually and semantically variable across images (Bainbridge et al., 2013). Therefore, it may be more difficult to identify the features of a scene that determine its memorability, causing incorrect memorability judgments. Conversely, it is possible that this variability across scenes ultimately helped participants improve their memorability judgments to a greater extent than participants viewing faces. However, due to the intrinsic differences between the face and scene stimuli in our study, these explanations are largely speculative.

Although participant memorability judgments were fairly accurate and improved over time, judgments were limited to highly memorable and forgettable images and the observed learning effect was small. Furthermore, final accuracy was still lower than that of a recent DNN’s predictions of the same scene images (Needell & Bainbridge, 2022). This implies a limit to the performance of human participants, who may be unable to access the full variance of memorability, or all the features that ResMem can access (Zhao et al., 2023). As ResMem is a “black box”, the exact dimensions it uses to generate predictions are still unknown, and thus it is difficult to directly compare human and DNN approaches. However, the only participant strategy resulting in significant learning—using characteristic features of the scene stimuli—parallels ResMem’s novel use of deep semantic information to aid its memorability estimation (Needell & Bainbridge, 2022).

Future steps could include explicitly instructing participants to adopt certain strategies (e.g., focusing on conceptual versus perceptual features), in order to boost accuracy of judgments and directly compare different influences on memorability. Additionally, testing participants’ own recognition memory for stimuli following training would allow for the comparison of memorability judgments with JOLs. Improving one’s understanding of image memorability could help teachers select memorable educational materials, students study forgettable test items, or researchers control for the memorability of their stimuli. Furthermore, these findings can offer insight into the lingering mysteries of memorability, metacognition, and our memories more broadly.

## Notes

### Competing Interest Statement

The authors have declared no competing interest.

