## Supplementary Material for "Learning Image Memorability with Feedback-Based Training"

301 Green Hall

5848 S. University Ave.

Chicago, IL 60637.

| Text/ Figure/ Table | Page |
| --- | --- |
| **Supplementary Text 1:** Main results excluding participants ≥ 65 years old. | 3 |
| **Supplementary Table 1:** Correlation results between face attribute ratings and guessed and true HR. | 4 |
| **Supplementary Figure 1:** Split-half consistency analysis of subjective memorability judgments. | 6 |
| **Supplementary Figure 2:** Correlation between guessed memorability and false alarm rate. | 7 |
| **Supplementary Table 2:** *T*-tests comparing Exp. 1 and 2 performance across time bins. | 8 |
| **Supplementary Figure 3:** Within-participant linear regression results of first and last half of trials. | 9 |
| **Supplementary Figure 4:** Within-participant linear regression results of replication and control experiments. | 11 |
| **Supplementary Table 3:** Mixed-effects logistic regression table. | 12 |
| **Supplementary Table 4:** Investigation of the interaction term from logistic regression results. | 13 |
| **Supplementary Figure 5:** Correlation between guessed and true memorability over time. | 14 |
| **Supplementary Table 5:** Examples of self-reported strategy categories. | 15 |
| **Supplementary Table 6:** Individual differences in task performance over time. | 17 |

**Supplementary Text 1**

In order to test the possibility that the learning, or lack of learning, of memorability may vary by age, we excluded all participants ≥ 65 years old and reran our main analyses to see if results replicated. This resulted in the exclusion of 3 participants from Exp. 1 and 5 participants from Exp. 2.

Results were qualitatively the same with older participants removed from analysis. As in the full results, the within-participant linear regression analysis showed no significant increase in accuracy in Exp. 1 (mean beta=0.010, SD=0.061); t(96)=1.56, $p>0.05$, but a significant increase in accuracy in Exp. 2 (M=0.017, SD=0.064); t(94)=2.52, $p=0.014$. Our mixed-effects logistic regression model resulted in an adjusted $R^2$ of 0.11 for Exp. 1, in which degree of memorability was predictive of accuracy (β=0.23, $p<0.001$), trial number was non-significant (β=0.03, $p>0.05$), and the interaction term between predictors was also non-significant (β=-0.005, $p>0.05$). For Exp. 2, the model resulted in an adjusted $R^2$ of 0.14, with trial number being significantly predictive of accuracy (β=0.04, $p=0.021$), degree of memorability being non-significant (β=-0.03, $p>0.05$), and the interaction term being significant (β=0.04, $p=0.014$). The significant interaction predictor was the only difference from the original results, in which the interaction term was marginally significant ($p=0.054$). Thus, we can conclude that the inclusion of the few participants ≥ 65 years old did not have a sizable impact on our main results.

**Supplementary Table 1**

*Correlation results between face attribute ratings and guessed and true HR*

| Face attribute | Guessed HR rho | Guessed HR *p* | True HR rho | True HR *p* |
|---|---|---|---|---|
| **common**/uncommon | -0.61 | 4.14 E-20 | -0.46 | 5.74 E-11 |
| **typical**/atypical | -0.52 | 4.98 E-14 | -0.44 | 6.48 E-10 |
| **memorable**/forgettable | 0.51 | 1.59 E-13 | 0.40 | 3.80 E-08 |
| **responsible**/irresponsible | -0.37 | 2.61 E-07 | -0.36 | 6.02 E-07 |
| **interesting**/boring | 0.35 | 1.43 E-06 | 0.19 | 0.01 |
| **familiar**/unfamiliar | -0.34 | 2.68 E-06 | -0.28 | 1.17 E-04 |
| **humble**/egotistic | -0.34 | 3.33 E-06 | -0.23 | 0.002 |
| **calm**/aggressive | -0.32 | 1.24 E-05 | -0.28 | 1.61 E-04 |
| **trustworthy**/untrustworthy | -0.31 | 2.17 E-05 | -0.26 | 5.30 E-04 |
| **normal**/weird | -0.30 | 4.11 E-05 | -0.31 | 3.08 E-05 |
| **emotionally stable**/unstable | -0.30 | 5.59 E-05 | -0.28 | 1.20 E-04 |
| **intelligent**/unintelligent | -0.26 | 3.77 E-04 | -0.29 | 5.83 E-05 |
| **kind**/mean | -0.26 | 3.62 E-04 | -0.22 | 0.004 |
| **caring**/cold | -0.24 | 0.002 | -0.22 | 0.003 |
| **friendly**/unfriendly | -0.17 | 0.02 | -0.18 | 0.02 |
| **emotional**/unemotional | 0.09 | 0.23 | 0.06 | 0.44 |
| **happy**/unhappy | -0.07 | 0.34 | -0.14 | 0.06 |
| **sociable**/introverted | 0.05 | 0.48 | -0.07 | 0.36 |
| **confident**/uncertain | 0.04 | 0.56 | -0.12 | 0.10 |
| **attractive**/unattractive | 0.03 | 0.71 | -0.05 | 0.49 |

*Note.* All face images used in Experiment 1 have crowdsourced ratings of 20 psychology and memory-related attributes and their antonyms (Bainbridge et al., 2013). We first confirmed that the ratings of all antonym pairs were significantly negatively correlated with each other across stimuli ($p < 10^{-4}$). We then averaged ratings of each attribute with those of its antonym (after subtracting the antonym from 10 to convert it to the same scale). A Spearman rank correlation was then calculated between ratings for each positive attribute (in bold) with guessed HRs and true HRs of Exp. 1 stimuli, using a false discovery rate (FDR) procedure to correct for multiple comparisons ($q < 0.05$). Rho and *p*-values are reported. Colored cells indicate FDR-corrected significance; green cells indicate a positive correlation and red cells indicate a negative correlation with the attribute in bold. As can be seen, associations of attributes with guessed and true HR were qualitatively identical. No correlation was greater than that of guessed and true memorability for faces (all $p>0.05$, confirmed by 10,000-iteration permutation test). This suggests that participants did not rely completely on any other attribute when making their memorability judgments.

**Supplementary Figure 1**

*Split-half consistency analysis of subjective memorability judgments*

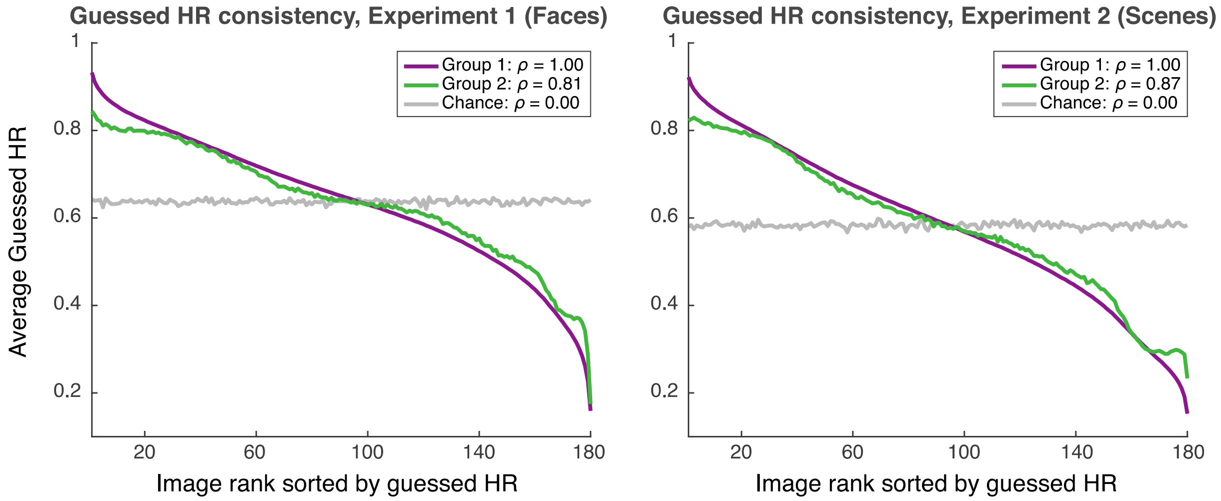


*Note.* Over 1000 iterations, the participant pool was randomly split into two halves, and a Spearman rank correlation was measured between the subjective memorability ratings of both groups. Purple lines correspond to the average guessed HR of each image (y-axis), sorted from highest to lowest (x-axis), according to group 1. Green lines correspond to the average guessed HR of the same images according to group 2, but again sorted by group 1's ranking. Gray lines correspond to chance-level memorability judgments, obtained by shuffling group 2's ranking. The average correlation between random halves of participants was high in both Experiment 1 (*left*; $\rho = 0.81$) and Experiment 2 (*right*; $\rho = 0.87$, both $p < 0.001$), as shown by the purple and green lines falling very close to each other. Thus, participants in both experiments tended to agree on the images they judged as memorable and forgettable.

**Supplementary Figure 2**

*Correlation between guessed memorability and false alarm rate*

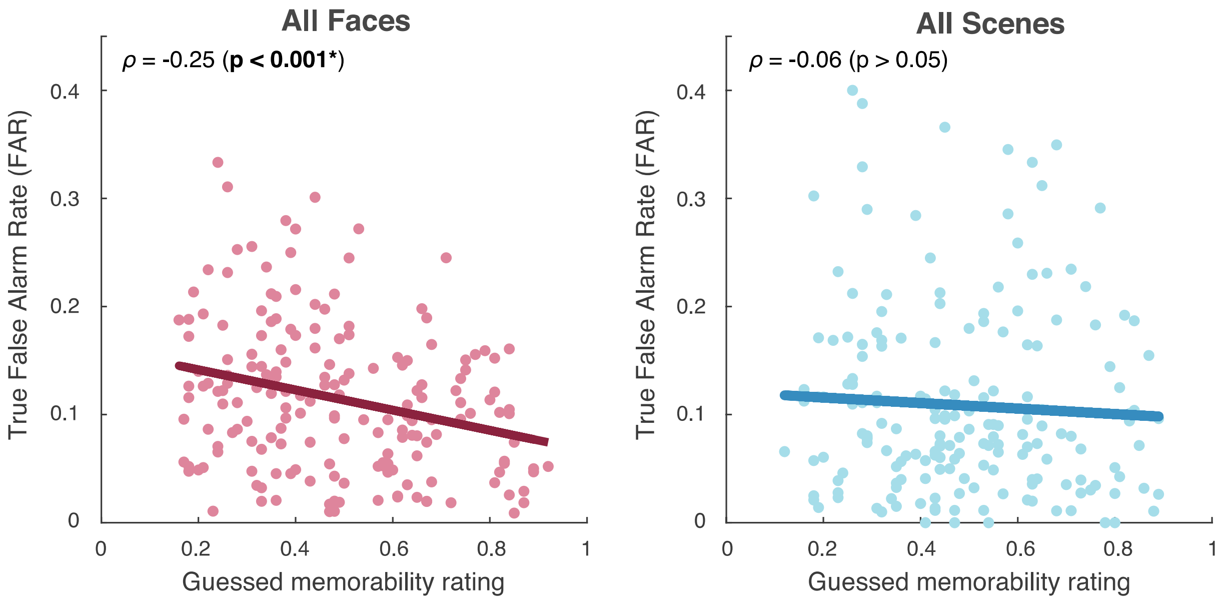


*Note.* Average guessed memorability rating is plotted along the x-axis and stimulus false alarm rate (FAR) is plotted along the y-axis for individual stimuli, represented by points in the scatterplots. For faces, there was a significant negative correlation between guessed memorability and FAR; in other words, participants assigned higher memorability scores to faces with lower false alarm rates. For scenes, there was no significant correlation between guessed memorability rating and FAR.

**Supplementary Table 2**

*T-tests comparing Exp. 1 and 2 performance across time bins*

| Bin number | Faces: mean % | Faces: SD of % | Scenes: mean % | Scenes: SD of % | *t*-statistic | *p*-value |
|---|---|---|---|---|---|---|
| 1 | 61.17 | 13.60 | 56.72 | 12.39 | 2.42 | 0.02 |
| 2 | 63.44 | 13.15 | 57.06 | 14.19 | 3.30 | 0.001 |
| 3 | 63.72 | 12.40 | 58.17 | 11.67 | 3.26 | 0.001 |
| 4 | 64.89 | 12.41 | 57.00 | 11.10 | 4.74 | 4.10 E-06 |
| 5 | 64.28 | 13.75 | 58.39 | 13.98 | 3.00 | 0.003 |
| 6 | 63.72 | 12.91 | 58.89 | 13.65 | 2.57 | 0.01 |
| 7 | 62.17 | 13.42 | 59.44 | 12.45 | 1.49 | 0.14 |
| 8 | 64.44 | 14.30 | 59.67 | 13.98 | 2.39 | 0.02 |
| 9 | 64.78 | 11.66 | 58.61 | 13.39 | 3.47 | 6.32 E-04 |
| 10 | 64.28 | 12.34 | 59.44 | 15.86 | 2.41 | 0.02 |

*Note*. Results of two-sample t-tests comparing average accuracy between Exp. 1 and 2 in each of 10 time bins, using an FDR procedure to correct for multiple comparisons ($q < 0.05$). For each time bin, average percent correct and standard deviation of percent correct in Exp. 1 and 2 are reported, as well as the *t*-statistic and *p*-value of the t-test comparing accuracy in both experiments. Green colored cells indicate bins in which accuracy for faces was significantly greater than accuracy for scenes, passing FDR correction.

**Supplementary Figure 3**

*Within-participant linear regression for first and last half of trials*

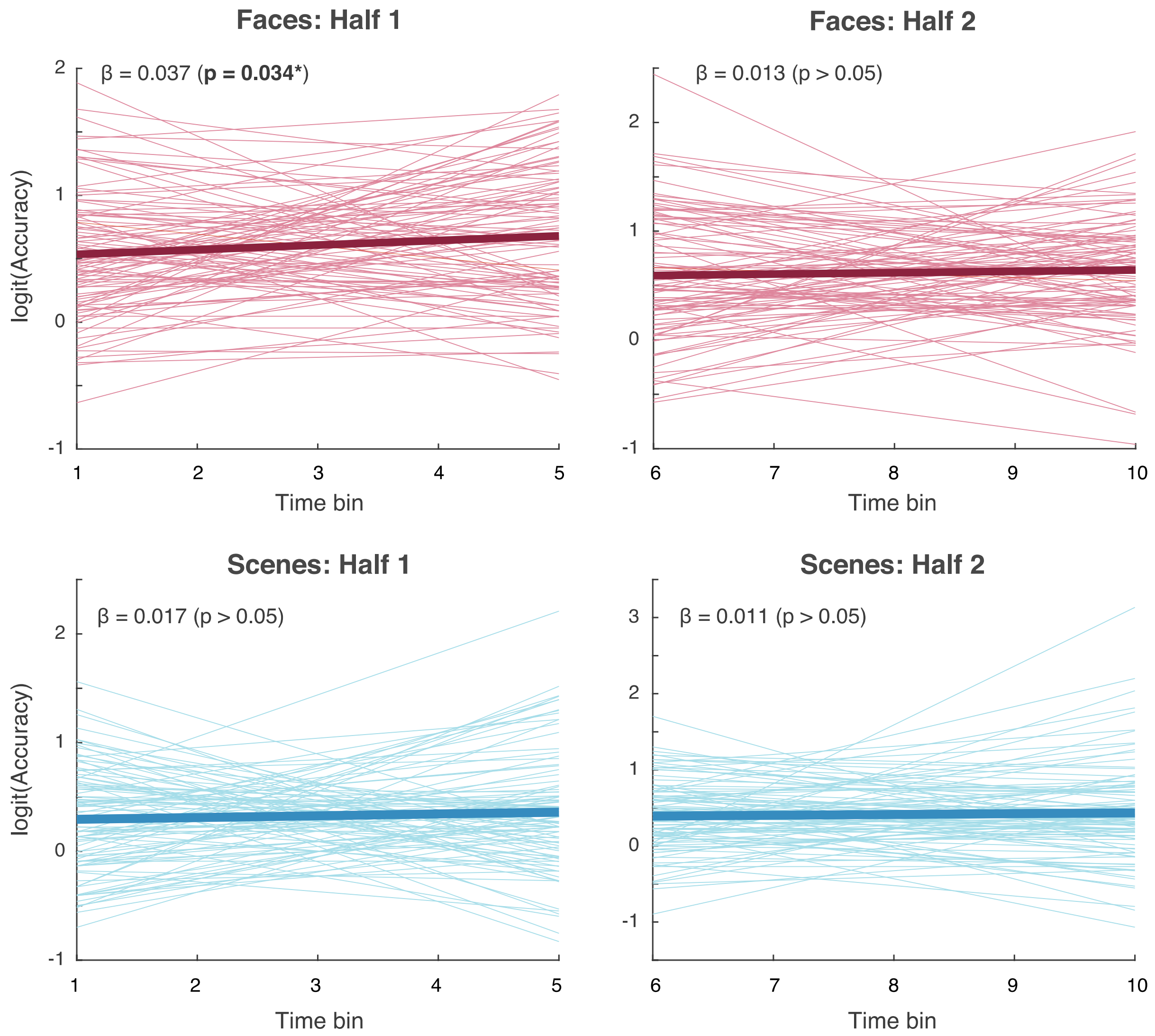


*Note*. We repeated the linear regression analysis as described in the main text, separately on the first half (time bins 1-5) and last half (time bins 6-10) of trials. As in Figure 4a-b, each line corresponds to a linear regression predicting logit-transformed accuracy (y-axis) from time (x-axis) for a single participant, and bolded lines correspond to the average regression lines. In the first half of Exp. 1, regression coefficients were significantly greater than zero, while in the second half, they were not significantly different, suggesting that perhaps participants were learning memorability early on in the task. However, when we directly compared coefficients in the first and second half, there was no significant difference ($p > 0.05$). For Exp. 2, neither half showed a significant difference from zero, and half 1 and 2 were also not significantly different from each other ($p > 0.05$).

**Supplementary Figure 4**

*Within-participant linear regression results of replication and control experiments*

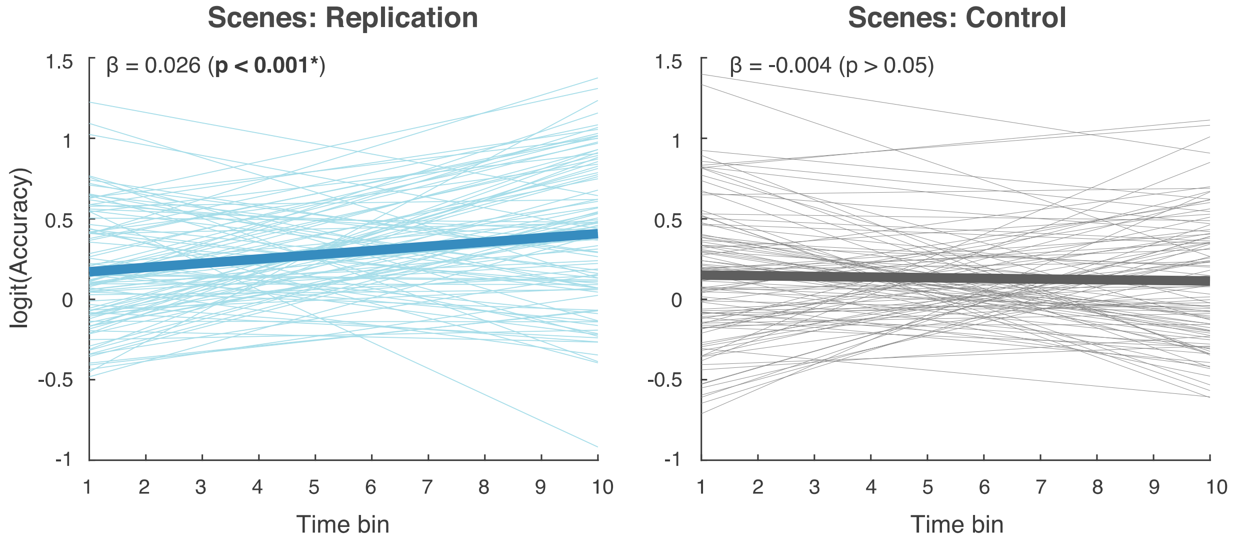


*Note.* Results from the replication experiment (left), with identical procedure to Experiment 2, as well as the no-feedback control experiment (right). As in Figure 4a-b, each line corresponds to a linear regression predicting logit-transformed accuracy (y-axis) from time (x-axis) for a single participant, and bolded lines correspond to the average regression lines. Regression coefficients were significantly greater than zero in the replication experiment, suggesting that, as in Exp. 2, participants learned memorability over time. Average accuracy was 53.6% (SD=15.1%) in time bin 1 and 59.3% (SD=13.1%) in time bin 10. In the control experiment, however, coefficients were not significantly different from zero—thus, participants showed no improvement in the absence of feedback. In this experiment, average accuracy was 53.1% (SD=14.2%) in time bin 1 and 52.3% (SD=13.6%) in time bin 10.

**Supplementary Table 3**

*Mixed-effects logistic regression table*

| Predictor | *β* | 95% CI | *p*-value | Odds ratio |
|---|---|---|---|---|
| | | Experiment 1: Faces | | |
| Trial number | 0.02 | [-0.008, 0.057] | 0.14 | 1.025 |
| Deg. of memorability | 0.23 | [0.132, 0.328] | 3.88 E-06 | 1.258 |
| Trial num:Deg. of mem. | -0.01 | [-0.042, 0.023] | 0.57 | 0.991 |
| | | Experiment 2: Scenes | | |
| Trial number | 0.04 | [0.007, 0.071] | 0.02 | 1.039 |
| Deg of memorability | -0.02 | [-0.141, 0.095] | 0.70 | 0.977 |
| Trial num:Deg. of mem. | 0.03 | [-0.001, 0.064] | 0.054 | 1.032 |

*Note.* Results of the mixed-effects logistic regression model predicting accuracy from trial number and degree of memorability, along with the interaction term (see main text). For each predictor, the beta estimate, 95% confidence interval of this estimate, *p*-value, and odds ratio are reported. In Exp. 1, degree of memorability was predictive of accuracy (significance indicated by the green cell). In Exp. 2, trial number was predictive of accuracy, and the interaction between predictors was marginally significant (indicated by the lighter green cell).

**Supplementary Table 4**

*Investigation of the interaction term from logistic regression results*

| **Degree of memorability quartile** | **Trial $\beta$** | **$\beta$ $p$-value** | **Adjusted $R^2$** |
|---|---|---|---|
| 1st quartile | -0.02 | 0.60 | 0.21 |
| 2nd quartile | 0.06 | 0.07 | 0.15 |
| 3rd quartile | 0.01 | 0.81 | 0.19 |
| 4th quartile | 0.12 | 3.67 E-04 | 0.21 |

*Note.* In the mixed-effects regression model, we included an interaction between the *trial number* and *degree of memorability* predictors, which was marginally significant in Experiment 2 ($\beta$ = 0.03, $p$ = 0.054). To investigate the nature of this interaction, we examined the effect of trial number on accuracy, separately over four quartiles of the degree of memorability measure. In more detail, quartile 1 included Exp. 2 stimuli with the lowest distance from 0.5 HR, and quartile 4 included stimuli with the greatest distance from 0.5 HR (the most memorable/most forgettable stimuli). For each of these quartiles, we fit a logistic regression model predicting accuracy from z-scored trial number, with participant and image as random intercepts. Trial beta estimate and *p*-value as well as model adjusted $R^2$ are reported. For stimuli in the 1st – 3rd quartiles of degree of memorability, trial number was not significantly predictive of accuracy. Trial number was only predictive of accuracy for scenes in the 4th quartile (significance indicated by the green cell). Thus, participants improved in accuracy over time primarily for the stimuli with the highest and lowest memorability scores.

**Supplementary Figure 5**

*Correlation between guessed and true memorability over time*

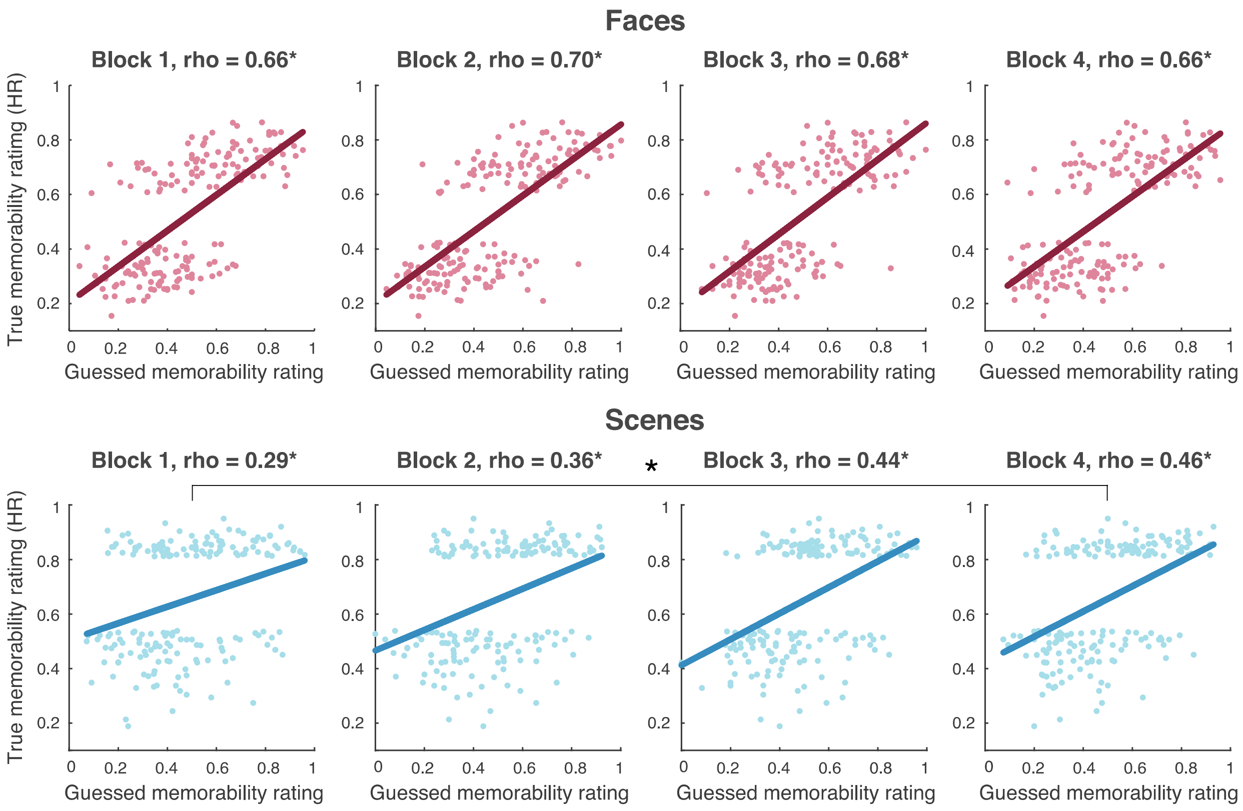


*Note.* As in Figure 2a, guessed memorability rating is plotted along the x-axis and true HR is plotted along the y-axis for all stimuli, represented by points in the scatterplots. Here, however, average guessed HR was calculated separately within four consecutive blocks of trials (block 1: trials 1-45, block 2: trials 46-90, block 3: trials 91-135, block 4: trials 136-180). For faces (*top*), there was no significant difference between correlations of guessed and true HR in any of the four time blocks ($p$>0.05; confirmed by 10,000-iteration permutation test). For scenes (*bottom*), this correlation was significantly different between blocks 1 and 4 ($p$<0.001; confirmed by 10,000-iteration permutation test).

* $p < 0.001$.

**Supplementary Table 5**

*Examples of self-reported strategy categories*

| **Strategy** | **Response examples** |
|---|---|
| Experiment 1: Faces | |
| Distinguishing feature(s) | - “Memorable faces had more distinguishing features”<br>- “Facial features like bright eyes, bangs, mustaches etc.”<br>- “Some memorable faces had earrings or piercings” |
| General face traits | - “Maybe photogenic vs not photogenic”<br>- “It seemed like a lot of the forgettable ones were middle-aged”<br>- “If their face is symmetrical and they’re good-looking, it’s hard to forget that face” |
| Expression | - “Strong expressions vs more neutral expressions”<br>- “People who looked very kind or super stoic”<br>- “I analyzed if they were smiling or not” |
| Thought would remember | - “I tried to think about what aspect of their face, if any, would help me remember them”<br>- “If I thought I would remember the persons face myself”<br>- “I looked at the face and asked myself if I see that person now, will I remember them tomorrow” |
| Gut feeling | - “Went with my gut decision”<br>- “Mostly just went with my own feelings” |
| Caught attention | - “I looked for something that caught my attention”<br>- “I thought if they stood out more then I chose memorable” |
| Image quality | - “High contrast versus low contrast faces”<br>- “The quality of the photo” |
| Using feedback | - “Trying to see the similarities of the image shown with those of the previous ‘memorable’ faces”<br>- “I tried to detect any sort of pattern to why someone was memorable or not” |

| Experiment 2: Scenes | |
|---|---|
| Specific objects/ places | - “Whether or not there is something in the picture that was usual or if it contained something that was out of the ordinary”<br>- “I would see if there was anything unique in the picture (i.e., a bathroom, a football field, a unique landscape)”<br>- “Were there specific things likes signs, or natural wonders” |
| General scene traits | - “Scenic pictures most of the time were not memorable”<br>- “Sometimes the beauty and composition of the image”<br>- “Symmetry of the photo” |
| Using feedback | - “I tried to compare each image against previous ones I'd seen”<br>- “I tried to remember what had been classified as what”<br>- “Based on what I got wrong” |
| Color | - “If the picture had…specific colors then it was memorable”<br>- “The vibrancy of the colors”<br>- “Bold colors made me choose memorable” |
| Gut feeling | - “I just followed my intuition”<br>- “Eventually it just became gut reaction” |
| Image quality | - "I thought it had something to do with…sharpness of the image”<br>- “The memorable pictures were clearer in focus” |
| Thought would remember | - “Just thought about if I would remember the image”<br>- “I just thought to myself: Will I be able to identify this picture if it appears once again?” |

*Note.* Examples of responses to the post-task questions about the perceived difference between memorable and forgettable images, and the strategies participants used to perform the task. Responses were grouped into different categories that were shared by at least 10 participants.

**Supplementary Table 6**

*Individual differences in task performance over time*

| Strategy category | *n* | Mean *β* | *β* SD | *t*-statistic | *p*-value |
|---|---|---|---|---|---|
| | | | Experiment 1: Faces | | |
| *Individual strategies* | | | | | |
| Distinguishing feature(s) | 80 | 0.012 | 0.063 | 1.75 | 0.08 |
| General face traits | 46 | 0.007 | 0.061 | 0.79 | 0.44 |
| Expression | 17 | 0.020 | 0.052 | 1.61 | 0.13 |
| Thought would remember | 16 | 0.009 | 0.060 | 0.58 | 0.57 |
| Gut feeling | 15 | 0.001 | 0.065 | 0.09 | 0.93 |
| Caught attention | 11 | 0.008 | 0.076 | 0.35 | 0.73 |
| Image quality | 11 | 0.007 | 0.073 | 0.32 | 0.75 |
| Using feedback | 10 | 0.013 | 0.043 | 0.97 | 0.36 |
| *Joint strategies* | | | | | |
| Dist. feat(s) & Gen. traits | 40 | 0.009 | 0.063 | 0.93 | 0.36 |
| Dist. feat(s) & Expression | 14 | 0.027 | 0.053 | 1.88 | 0.08 |
| Dist. feat(s) & Thought would remember | 11 | -0.003 | 0.067 | -0.16 | 0.88 |
| Dist. feat(s) & Gut feeling | 11 | 0.008 | 0.070 | 0.37 | 0.72 |
| Gen. traits & Expression | 11 | 0.017 | 0.046 | 1.24 | 0.24 |
| | | | Experiment 2: Scenes | | |
| *Individual strategies* | | | | | |
| Specific objects/ places | 63 | 0.029 | 0.068 | 3.34 | 0.001 |
| General scene traits | 32 | 0.004 | 0.039 | 0.52 | 0.61 |
| Using feedback | 20 | 0.032 | 0.067 | 2.15 | 0.044 |
| Color | 19 | 0.037 | 0.061 | 2.63 | 0.017 |
| Gut feeling | 14 | 0.004 | 0.054 | 0.29 | 0.78 |
| Image quality | 12 | -0.006 | 0.070 | -0.28 | 0.78 |
| Thought would remember | 10 | 0.003 | 0.063 | 0.13 | 0.90 |
| *Joint strategies* | | | | | |

| | | | | | |
|---|---|---|---|---|---|
| Spec. objects & Gen. traits | 16 | 0.004 | 0.043 | 0.34 | 0.74 |
| Spec. objects & Feedback | 15 | 0.040 | 0.068 | 2.28 | 0.039 |
| Spec. objects & Color | 18 | 0.041 | 0.059 | 2.94 | 0.009 |

*Note.* Participants were assigned to one or more category according to their self-reported strategies for completing the task. Individual categories containing 10 or more participants are listed here. Combinations of two categories containing 10 or more participants are also listed. Within each group, we ran the within-participant linear regression analysis to test for a significant change in logit-transformed accuracy over the full task (see main text). We used an FDR procedure to correct for multiple comparisons, separately for individual and joint strategy groups ($q < 0.05$). For each response category, the number of participants, mean and standard deviation of participants' regression coefficients, and *t*-statistic and *p*-value from the *t*-test on beta values are reported. Colored cells indicate significance; darker green cells indicate strategies that passed FDR correction and lighter green cells indicate strategies that failed FDR correction but showed a marginally significant effect ($p < 0.05$). No individual or joint strategies in Exp. 1 exhibited a significant effect. In Exp. 2, one individual strategy and one joint strategy resulted in a significant increase in accuracy over time, with the remaining strategies exhibiting marginal increases or no changes in performance.
